# Host related factors determine co-occurrence patterns between pathogenic bacteria, protozoa, and helminths in populations of the multimammate mouse, *Mastomys natalensis*

**DOI:** 10.1101/2022.01.14.476303

**Authors:** Joachim Mariën, Bram Vanden Broecke, Pamela Jones June Tafompa, Lisse Bernaerts, Alexis Ribas, Ladslaus L. Mnyone, Loth S. Mulugu, Herwig Leirs

## Abstract

Advances in experimental and theoretical work increasingly suggest that parasite interactions within a single host can affect the spread and severity of wildlife diseases. Yet empirical data to support predicted co-infection patterns are limited due to the practical challenges of gathering convincing data from animal populations and the stochastic nature of parasite transmission. Here, we investigated co-infection patterns between micro- (bacteria and protozoa) and macroparasites (gastrointestinal helminths) in natural populations of the multimammate mouse (*Mastomys natalensis*). Fieldwork was performed in Morogoro (Tanzania), where we trapped 211 individual *M. natalensis* and tested their behavior using a modified open-field arena. All animals were checked on the presence of helminths in their gastrointestinal tract, three bacteria (*Anaplasma, Bartonella*, and *Borrelia*) and two protozoan genera (*Piroplasma* and *Hepatozoon*). Besides the presence of eight different helminth genera (reported earlier), we found that 21% of *M. natalensis* were positive for *Anaplasma*, 13% for *Bartonella*, and 2% for *Hepatozoon* species. Hierarchical modelling of species communities was used to investigate the effect of the different host-related factors on these parasites’ infection probability and community structure. Our results show that the infection probability of *Anaplasma* and *Bartonella* was higher in adults than juveniles. We also observed that females and less explorative individuals had a higher infection probability with *Bartonella*. We found limited support for within-host interactions between micro-and macroparasites, as only animals infected with *Bartonella* were significantly more likely to be infected with *Protospirura, Trichuris*, and *Trichostrongylidae* helminths.

## Introduction

In natural populations, hosts are continuously exposed to multiple parasites while being acutely infected with others at the same time. Because a parasite can directly or indirectly affect the host’s susceptibility to another parasitic infection, co-infections may have important implications for the spread and pathogenicity of a disease (Clark et al., 2016; Henrichs et al., 2016; McArdle et al., 2018; Pedersen and Fenton, 2019). For example, immunodeficiency caused by chronic HIV infections increases the severity of several other human diseases, such as tuberculosis, cryptococcosis, hepatitis B, hepatitis C virus, and malaria (McArdle et al., 2018). Direct interactions between parasites can occur due to resource competition or predation of one parasite on the other (Pedersen and Fenton, 2019; Vaumourin et al., 2015). Indirect interactions are often mediated through the host’s immune response and can be facilitative or inhibitory (Ramsay and Rohr, 2021). For instance, microparasites generally trigger the T helper 1 (Th1) branch of the adaptive immune response, whereas macroparasites trigger the T helper 2 (Th2) branch (Blanco and Garcia, 2008; McArdle et al., 2018). By activating one of these two branches, the host’s resources will become depleted so much that it cannot properly ward off another infection. Hosts are therefore assumed to be more susceptible to microparasite infections after infection with macroparasites. Indeed, African buffalos have a weaker Th1 response after nematode infections, facilitating the invasion of bovine tuberculosis (Ezenwa et al., 2010). In contrast, co-infections between two micro- or two macroparasites are more likely to be inhibitory because of cross-immunity and the activation of the same T helper immune response (Ramsay and Rohr, 2021).

Studying rodent-borne parasites improved our understanding of the importance of co-infections over the past decades (Pedersen and Fenton, 2019). While laboratory experiments typically inoculated mice with controlled doses of two or more parasites, field experiments allowed to investigate the importance of co-infections under natural conditions. Those field studies mainly focused on micro-micro or macro-macroparasite interactions (Pedersen and Fenton 2019). For example, one of the most convincing studies found that field voles (*Microtus agrestis*) were more likely to be infected with *Anaplasma, Babesia*, and *Bartonella* bacteria after a cowpox virus infection, but less likely to be infected with *Bartonella* after a *Babesia* infection (Telfer et al., 2010). Macro-microparasites interactions are less studied in rodent populations (Pedersen and Fenton 2019). One study found that bank voles (*Myodes glareolus*) infected with the nematode *Heligmosomoides mixtum* were more likely to have antibodies against Puumala hantavirus, which was explained by the immunomodulating effects of the helminth infection on the Th1 immune response (Salvador et al., 2011). In contrast, microparasites had no significant impact on the presence of the nematode *Heligmosomoides polygyrus*, suggesting that direct micro-macroparasite interactions are relatively rare in natural conditions (Luong et al 2010).

This study focuses on micro-macroparasite interactions in the multimammate mouse (*Mastomys natalensis*), the most important rodent pest species in sub-Saharan Africa (Leirs et al., 1990; Mulungu, 2017). While its natural habitat consists of savannah and grassland areas, the animal currently thrives in agricultural fields and human dwellings where outbreaks can cause crop losses up to 80% (Mulungu, 2017; Mwanjabe et al., 2002). The rodent also hosts several zoonotic diseases, including *Yersinia pestis* (bubonic plague), *Leptospira interrogans* (leptospirosis), *Leishmania donovani* (cutaneous leishmaniasis) and Lassa virus (Lassa fever) (Holt et al., 2006; Mariën et al., 2019a; Meerburg et al., 2009; Monath, 1987; Neerinckx et al., 2008; Sadlova et al., 2019) as well as several ecto- and endoparasites (Brouat et al., 2007; Brouat and Duplantier, 2007; Diagne et al., 2016; Diouf et al., 2013; Oguge et al., 1997; Ribas et al., 2017, 2013, 2012). We recently used *M. natalensis* as a model system to investigate how host characteristics and behaviour affect the community structure of gastrointestinal helminths (Vanden Broecke et al., 2021a). We observed that helminth richness was higher in adults and females than juveniles and males. Additionally, we found that less explorative individuals (observed in a modified open field arena) had a higher infection probability with different helminths. Here, we screened the same individuals on the presence of five microparasite genera known to cause disease in humans and livestock: *Bartonella*, *Anaplasma, Borrelia, Piroplasma*, and *Hepatozoon. We* hypothesize that the host’s gastrointestinal helminth community has a significant effect on the presence of these microparasites due to immunomodulation. More specifically, we predict that infection with helminths will increase the probability of becoming infected with these microparasites, even after correcting for host characteristics such as sex and age. Additionally, we expect that highly explorative individuals are more likely to be infected with microparasites compared to less explorative individuals due to the higher chances of encountering an infected vector (e.g., tick, lice, mites) throughout their life (Barber and Dingemanse, 2010; Bohn et al., 2017; Boyer et al., 2010).

## Material and methods

### 2.1. Study site and trapping

The field setup, rodent collections, behavioral measurements, and helminth screening are described in detail in Vanden Broecke et al. (2021a). In brief: rodents were trapped on eleven different sites in both maize fields and in fallow lands on the Sokoine University of Agriculture farm (SUA) in Morogoro, Tanzania, from July until September 2019. Traps were placed in the late afternoon and checked in the early morning, and captured rodents were brought to the Institute of Pest Management Centre (IPM).

The behaviour of the rodents was measured immediately after they arrived at the IPM using a hole-board test, which is commonly used to study behaviour in *M. natalensis* (Vanden Broecke et al., 2021b, 2021c, 2021a, 2019). Behavioral recordings started when the individual was inside the box and lasted for 10 minutes. During this period, we measured five different behaviors: activity, the number of times they sniffed one of the blind holes, number of head dips, the time they spent grooming and the number of jumps. After each recording, the box was cleaned with 70% ethanol to remove animal scent and dirt. For all animals, we noted their body weight, sex, and reproductive status, following Leirs et al. (1994). Blood samples were taken from the retro-orbital sinus when the animal was still alive and was preserved on pre-punched filter paper (Serobuvard, LDA 22, Zoopole). We then euthanized the rodents using a halothane overdose followed by cervical dislocation. We collected a small piece of the kidney, liver, lung, salivary glands, and brain and stored it in 100% ethanol. Afterward, we removed the whole gastrointestinal tract and kept it in a 50 ml tube with 100% ethanol for further analysis at the parasitology lab at the University of Barcelona (Ribas et al., 2011). The helminths were identified to genus or species level using morphological characteristics described in (Ribas et al., 2017, 2013; Vanden Broecke et al., 2021a).

### 2.2. Pathogen DNA detection

Genomic DNA from the tissues was extracted using the NucleoSpin^®^ Tissue DNA Extraction kit (MACHEREY-NAGEL GmbH & Co. KG, Germany) according to the manufacturer’s protocol. We pooled tissues of the kidney, liver, and lungs per individual to have 25 mg in total. To ensure no contamination when handling specimens from different animals, the knife, scissors, and tweezers were decontaminated using 5 % Virkon^®^ (Antec International) and dried after collection from each rodent. DNA quality and quantity were checked for each sample using a Qubit fluorometer from Thermo fisher Scientific. The DNA extracts were then stored at −20 °C until PCR analysis.

First, all DNA extracts were screened on the presence of *Bartonella, Anaplasma, Borrelia*, and *Piroplasma* using real-time qPCR systems with primers and probes that have a width specificity (Dahmana et al., 2020). Amplification reactions were conducted in a final volume of 20 μL containing 10 μL of 2xEurogentec Takyon^™^ Mix (Eurogentec, Liège, Belgium), 1 μL of each primer (0.5 μM), 2.5 μL of DNase-free water, and 5 μL of DNA template. The RT-qPCR was performed on the StepOne^™^ Real-Time PCR system (by Thermo Fisher Scientific) using the following thermal profile: an incubation step at 50 °C for two minutes for eliminating PCR amplicons’ contaminant, then an activation step at 95 °C for three minutes followed by 40 cycles of denaturation at 95 °C for 15 seconds and an annealing-extension at 60 °C for 30 seconds. Samples with a Ct value below 35 were screened a second time (duplicates). An individual was considered positive on the RT-qPCR if tested positive during both runs.

We then screened all extracts on the presence of *Hepatozoon* using a broad-range conventional PCR system (Dahmana et al., 2020). We also used conventional PCRs to amplify and sequence all qPCR-positive samples for *Anaplasma* targeting the 23S gene (Dahmana et al., 2020) and *Bartonella* targeting the gltA and 16S-23S rRNA ITS region (Böge et al., 2021). The amplification reactions were conducted in a final volume of 25 μL, containing 12.5 μL of Amplitaq Gold master mix, 0.5 μL of each primer, 2.5 μL of DNA template, and 9 μL of DNA free water. Reactions were performed in an automated thermal cycler (TProfessional Basic Thermocycler by Biometra) following the specific thermal cycling profile. For A*naplasma* and *Hepatozoon:* one incubation step at 95 °C for 15 min, 40 cycles of 60sec at 95 °C, 30sec at annealing temperature and 1min at 72 °C and a final extension step of 5min at 72 °C. Amplification of the gltA *Bartonella* gene consisted of 45 cycles of denaturation at 95 °C for 30 s, annealing at 53 °C for 30 s and elongation at 72 °C for 1 min. Amplification of the ITS *Bartonella* gene consisted of 40 cycles for 30 s at 94 °C, for 30 s at 66 °C, for 50 s at 72 °C. PCR products were prepared with DNA Gel Loading Dye (6×) (Thermo Fisher Scientific Baltics UAB, Vilnius, Lithuania) for gel electrophoresis in 2% agarose. The results were visualized by UV light using the Syngene Geneflash Network Bio-Imaging device. Amplicons of positive samples were purified and sent to the Vlaams Institute of Biotechnologie for Sanger Sequencing with forward and reversed primers. Sequences were trimmed using Geneious and compared to available data in GenBank with BLASTn. Individuals were considered to be positive for *Anaplasma* or *Bartonella* if they were positive on the RT-qPCR after replication, and we obtained Sanger sequences from the conventional PCR for at least one gene. If no sequences were obtained, they were considered uncertain if we obtained at least repeated positive results on the RT-qPCR. The categorization was made because the RT-qPCR is generally a more sensitive approach than the conventional PCR (which targets larger DNA amplicons).

### 2.3. Statistical analysis

We used Hierarchical Modelling of Species Communities (HMSC) to test which host characteristics and behaviors affect the infection probability of *Bartonella* and *Anaplasma* and if co-infection patterns exist with the different helminths (Ovaskainen et al. 2017; Ovaskainen and Abrego 2020). We did not consider the other microparasites because they were either not found (*Borrelia* and *Piroplasma*) or were present at a too low prevalence (*Hepatozoon* and *Ehrlichia*; see results). We used the same models and input data as described in Vanden Broecke et al. (2021a) but added the presence of *Bartonella* and *Anaplasma* as additional response variables in the response matrix (Ovaskainen et al. 2017; Ovaskainen and Abrego 2020). We used a presence-absence model in which we transformed the response variable to a binomial variable where the parasite was either present (1) or absent (0) within a specific individual (Ovaskainen and Abrego 2020). To correct for the presence of uncertain samples (i.e., individuals that were repeatedly positive on the qPCR but negative for Sanger sequencing on the conventional PCR), we ran two different models where we considered all uncertain samples either as negative or positive (further referred to as, respectively, the uncertain-negative and uncertain-positive model). This allows testing if the assignment of the uncertain status affects the main results and conclusions.

We modeled the individuals’ identity as the sampling unit, which was nested as a random effect within the field site where the individual was trapped. As explanatory variables, we included the individuals’ sex (male or female), reproductive age (adult or juvenile), their exploration and stress-sensitivity behavior expressed in the hole-board test (more information in Vanden Broecke et al. 2019). Additionally, we included the results of antibody status against Morogoro arenavirus (MORVab), for which we screened the animals in the previous study. HMSC models were fitted with the R-package Hmsc (version 3.0-9; Tikhonov et al. 2020) using prior default distributions (Ovaskainen and Abrego, 2020). We sampled the posterior distribution with five Markov Chain Monte Carlo (MCMC) chains, each run with 3,000,000 iterations of which the first 1,000,000 were removed as burn-in. The chains were thinned by 1,000 to yield 2,000 posterior samples per chain, resulting in 10,000 posterior samples in total. We examined MCMC convergence using the potential scale reduction factors of the model parameters (Ovaskainen and Abrego, 2020). Both models’ explanatory and predictive power were analyzed using the species-specific AUC and Tjur’s R^2^ (Tjur, 2009) values (Pearce and Ferrier, 2000). Explanatory power was computed by making predictions based on models fitted to all data. Predictive ability was calculated by performing 5-fold cross-validation. The sampling units were assigned randomly to five-folds, and predictions for each fold were based on models fitted to data on the remaining four-folds.

## Results

### PCR analysis

We had a total sample size of 211 individuals (*N*_male adult_ = 54, *N*_male juvenile_ = 21, *N*_female adult_ = 112, *N*_female juvenile_ = 24). Overall 28 individuals tested positive for *Bartonella* (13.3% CI 9.0-18.6%). A complete sequence for the gltA and ITS genes was obtained for 14 and 15 individuals, respectively. For ten individuals, we did not obtain any sequence, despite they were repeatedly positive in the RT-qPCR (these individuals were given the uncertain *Bartonella* status). All sequences belonged to the same *Bartonella* strain (OL984911) with the highest similarity in Genbank to *Bartonella mastomydis* (KY555067, 95.53%) closely followed by *Bartonella elizabethae* (LR134527, 95.48%) based on both genes. For *Anaplamsa*, 44 individuals tested positive (20.9% CI 15.5-26.9%). A complete amplicon sequence of the 23S gene was obtained for 35 individuals. The sequences belonged to two different strains (OL982744, OL982748). One strain was most similar to an uncultured *Anaplasma* strain (MT269273, 96.9%, *n*=33) in Genbank; the other strain was most similar to an uncultured *Ehrlichia* strain (MK942592, 98.13%, *n*=2). We did not obtain any sequence for nine individuals, although they were repeatedly positive on the RT-qPCR (*Anaplasma* uncertain status). For *Hepatozoon*, five individuals tested positive (2.4% CI 1.0-5.4%), and all belonged to the same strain (OL982745), which was highly similar to *Hepatozoon ophisauri* (MN723845, 99.1%) in Genbank. All animals tested negative for *Borrelia* and *Piroplasma*.

### Hierarchical modelling of species

To investigate if co-infections between the micro and macroparasites occurred and to test if any host characteristics and behaviors could affect the infection probability of *Bartonella* and *Anaplasma*, we constructed two HMSC models in which we assigned the uncertain samples either as negative or positive. The MCMC convergence of the two models was satisfactory: the potential scale reduction factors for the *β*-parameters (which measured the responses of the different parasitic species to the other intrinsic covariates; Ovaskainen et al. 2017) was on average 1.006 (max = 1.103) for the negative model and 1.000 (max = 1.001) for the positive model. Both models fitted the data adequately, with a mean AUC of 0.868 (min 0.730 – max 0.996) for the explanatory power and 0.655 (0.510 – 0.792) for the predictive power in the uncertain-negative model (supplementary Figure 1A), and a mean AUC of 0.863 (0.718 – 0.992) for the explanatory power and 0.667 (0.505 – 0.804) for the predictive power in the positive model (supplementary Figure 2A). The mean Tjur R^2^ for the explanatory power was on average 0.209 (0.066 – 0.387) and 0.075 (0.003 – 0.231) for the predictive power for the negative model (supplementary Figure 1B), and the mean Tjur R2 for the explanatory power was on average 0.227 (0.065 – 0.422) and 0.092 (0.000 – 0.210) for the predictive power in the positive model (supplementary Figure 2B).

Variance partitioning showed that the hosts’ age explained a substantial proportion of the variation in the uncertain-negative (16.3%; supplementary Figure 3A) and positive (16.2%; supplementary Figure 3B) models. Indeed, both models show that the infection risk with *Bartonella* and *Anaplasma* is higher in adults compared to juveniles (Figure 1, supplementary Table 1). The hosts’ behaviour was responsible for 16.2% of the variation in the uncertain-negative model (exploration = 10.0%; stress-sensitivity = 6.2%; supplementary Figure 3A) and 14.2% in the uncertain-positive model (exploration = 9.4%; stress-sensitivity = 4.8%; supplementary Figure 3B). However, the individual’s behaviour had only an effect on infection risk with *Bartonella*, where we found a significant negative effect of both exploration and stress-sensitivity on the infection risk with *Bartonella* (Figure 2, supplementary table 1). The hosts’ sex was responsible for 6.4% of the variation in the uncertain-negative model (supplementary Figure 3A) and 6.7% in the uncertain-positive model (supplementary Figure 3A 3B). We did find that females were significantly more likely to be infected with *Bartonella* than males. This effect was only present in the uncertain-positive model and not in the uncertain-negative model (Figure 3, supplementary Table 1), which may suggest that the effect is not well supported. There were no differences between males and females in infection risk with *Anaplasma* (supplementary Table 1). The individual identity, which was included as a random effect in both models, explained the largest part of the variation in both models (negative model: 19.0%; positive model: 19.4%; supplementary Figure 3A), which suggests that a large proportion of the variance can be ascribed to unaccounted variation among the individuals.

**Figure 1:**
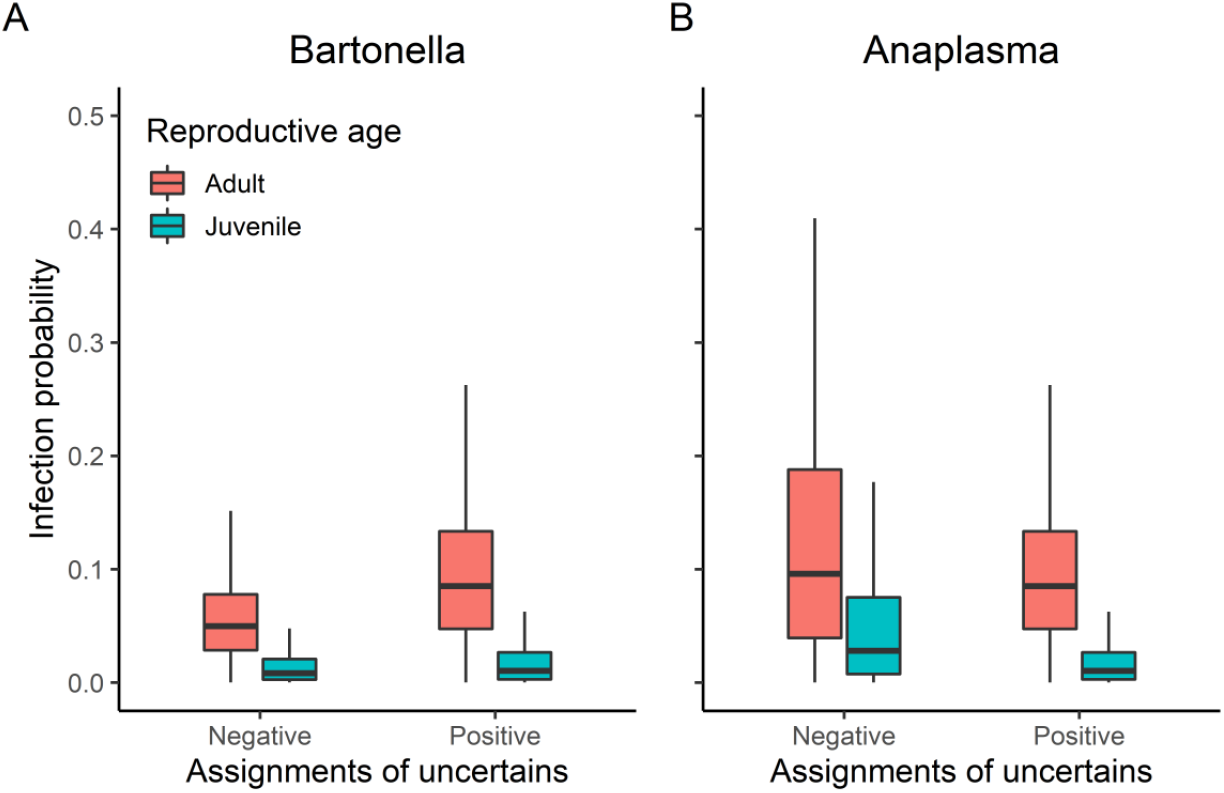
Predicted infection probability of *Mastomys natalensis* per reproductive age category for *Bartonella* **(A)** and *Anaplasma* species **(B)**. Different infection probabilities were estimated when considering uncertain samples to be positive or negative. Boxplots show the estimated infection probability of all posterior samples (*n*=10,000).

**Figure 2:**
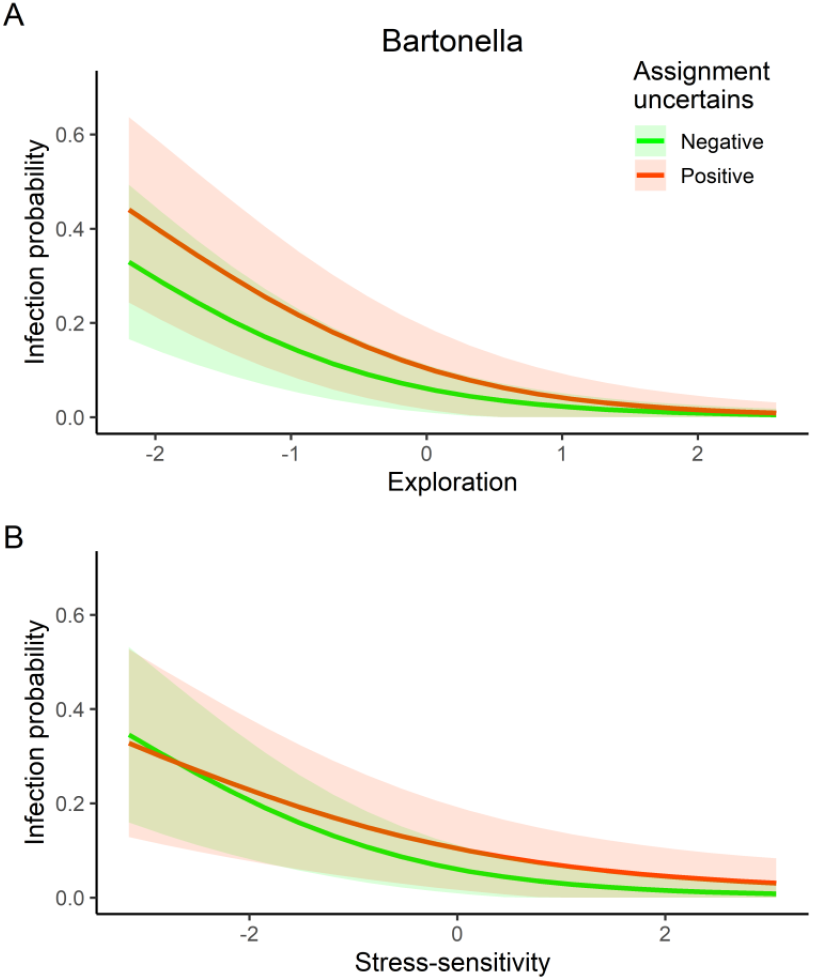
Predicted infection probability of *Mastomys natalensis* as a function of exploration **(A)** and stress-sensitivity **(B)**. Different infection probabilities were estimated when considering uncertain samples to be positive or negative. Envelops represent 95% credible intervals on the estimated infection probability of all posterior samples (*n*=10,000).

**Figure 3:**
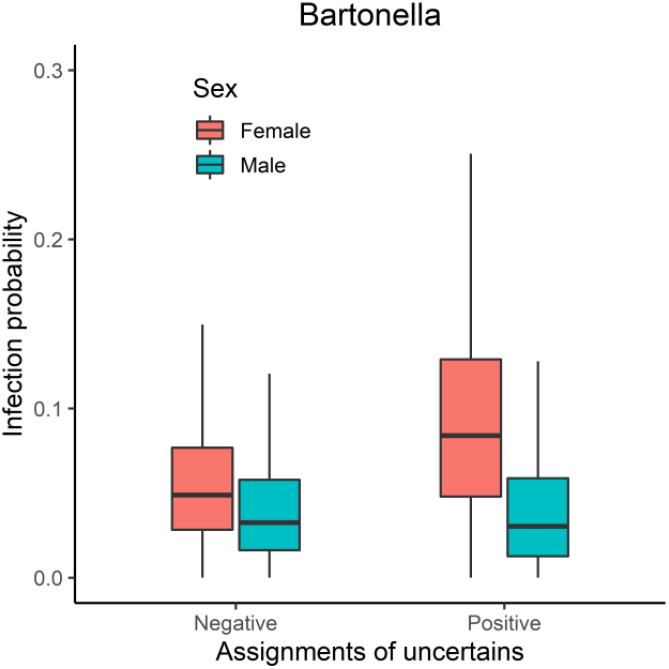
Predicted infection probability of *Mastomys natalensis* for *Bartonella* per sexual condition. Different infection probabilities were estimated when considering uncertain samples to be positive or negative. Boxplots show the estimated infection probability per 10,000 posterior samples.

The model showed a positive co-occurrence pattern between *Bartonella* and three helminths, namely: *Protospirura muricola*, *Trichuris mastomysi*, and *Trichostrongylidae* at the individual level after controlling for host-associated and spatial confounding factors (Figure 4 and 5A). This effect, however, was only signficant in the uncertain-positive model and not in the uncertain-negative model (Figure 4A,C and 5A), suggesting that this effect is not well supported. Finally, we found that the site in which the individuals were trapped explained a large proportion of the variation in both the uncertain-negative (16.4%, supplementary Figure 3A) and positive model (15.9%, %, supplementary Figure 3B), which suggests that local variation among the sites has a large effect on the presence of certain parasites. Indeed, we found that *Anaplasma* and *Hymenolepis* co-occurred more frequently together, at the site level, than expected by random in both models (Figure 4B,D and 5B). The co-occurrence of different parasites at the different sites was even stronger in the positive model, where we found a negative co-occurrence pattern between two groups (*Anaplasma – Hymenolepis* and *Davaineidae – Syphacia*; Figure 4D and 5B) which co-occurred positive with each other within each group. This suggests that rodents at a specific location are either infected with *Anaplasma-Hymenolepis* or with *Davaineidae-Syphacia*.

**Figure 4:**
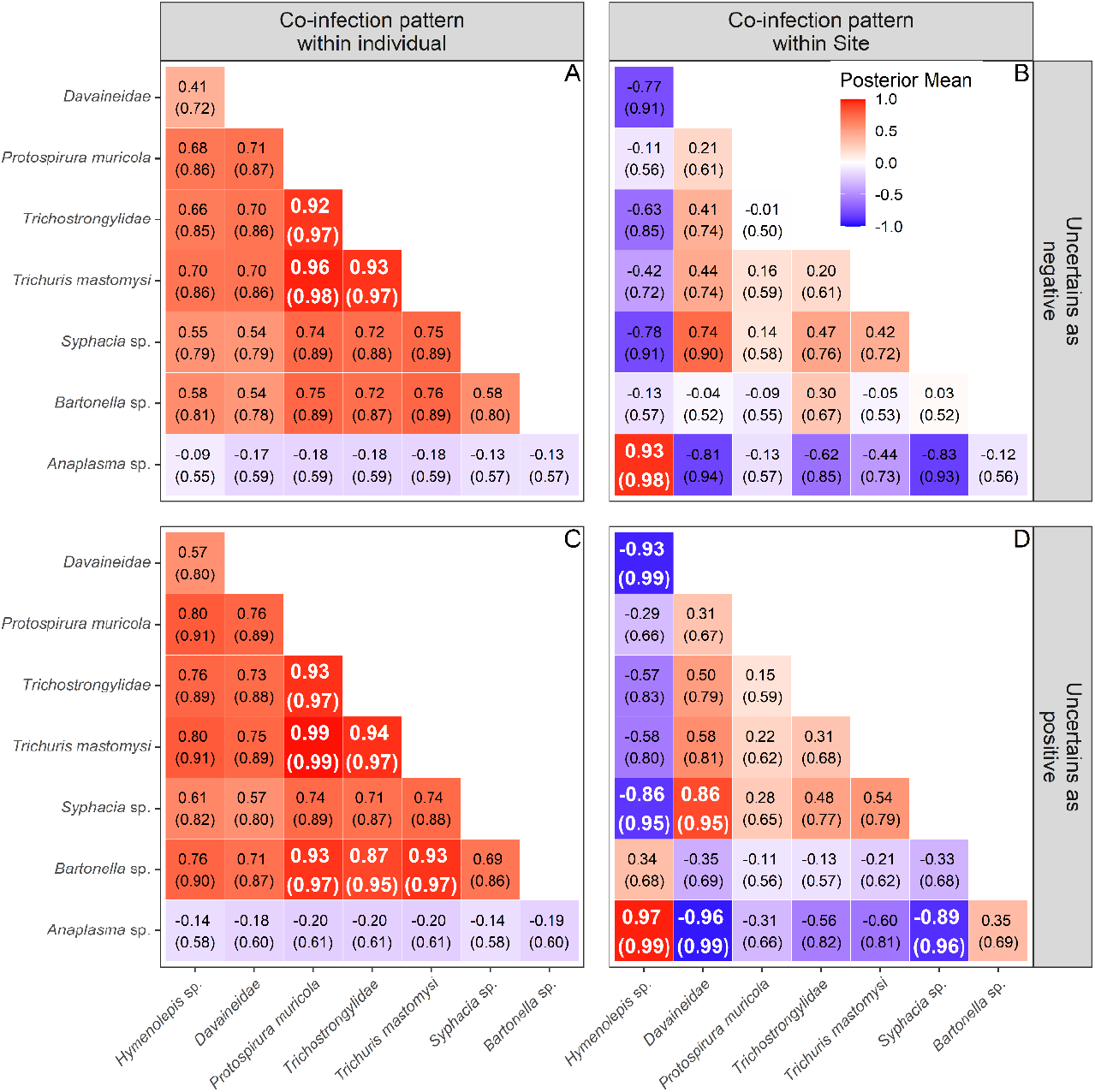
Co-infection patterns (posterior mean and support) of the different micro-and macroparasites detected in *Mastomys natalensis* based on the presence-absence HMSC model after controlling for host-associated factors (sex, age and behaviour). Different co-infection patterns were estimated for the individuals’ identity as the sampling unit and the field site where the individual was trapped. Different co-infection patterns were also assessed when considering uncertain samples to be positive or negative.

**Figure 5:**
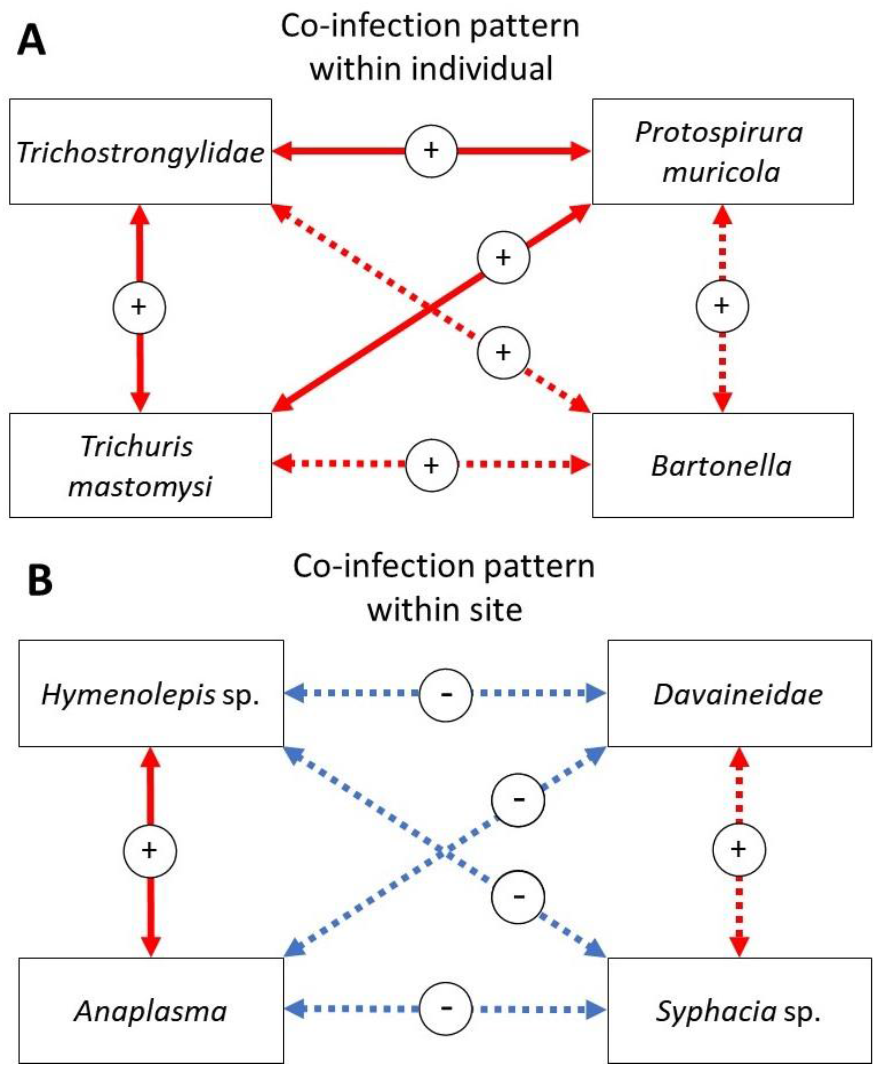
Graphical summary of the results from the HMSC model. Arrows indicate strong correlations, with 95% posterior probability support, between two parasites. Full arrows indicate strong correlations found in both models, while dashes arrows indicate strong correlations when considering the uncertain samples as positives. Red arrows indicate positive correlations and blue arrows negative.

## Discussion

In this study, we investigated the link between the gastrointestinal helminth community and the presence of different microparasites in *M. natalensis*. Three microparasites genera were detected from the five included in our screening panel: *Bartonella, Anaplasma*, and *Hepatozoon*, which is, to our knowledge, the first observation of these pathogens in rodent populations in Morogoro, Tanzania. We found that infection risk with *Bartonella* and *Anaplasma* was higher in adults. Additionally, we found that the host’s sex and behaviour affected *Bartonella* infection risk, but not *Anaplasma*. Lastly, our data suggests that individuals infected with *Bartonella* are more likely to be infected with different gastrointestinal helminths.

*Bartonella* is a gram-negative intracellular, flea-borne bacteria with a worldwide distribution. They infect erythrocytes, macrophages and endothelial cells of various mammals (Dehio, 2004). At least 13 *Bartonella* species are responsible for human diseases, including *B. bacilliformis*, *B. quintana* and *B. henselae* which causes, respectively, Carrion disease, trench fever and cat-scratch disease (Gutiérrez et al., 2015). The *Bartonella* species that we detected in *M. natalensis* in Morogoro was genetically most closely related to *B. mastomydis*, which was previously isolated from *Mastomys erythroleucus* in Senegal (Dahmani et al., 2018). The probability of being infected with *Bartonella* increased with age and was higher in females compared to male mice. The increase with age is expected given that *Bartonella* infections are usually long-lasting in the natural reservoirs (up to several months in European rodent populations), and older individuals have more chance of acquiring an infection during their life (Gutiérrez et al., 2015). Most studies reported no difference in *Bartonella* prevalence between sexes. However, minor differences were observed in *Apodemus sylvaticus* and *A. flavicollis*, in which males presented higher infection prevalence than female rodents (Gutiérrez et al., 2015; Welc-Faleciak et al., 2010). Our results suggest that female rodents are more likely to be infected with *Bartonella* for which there are two potential explanations. A first explanation results from the unusual social behavior (non-territorial) of female *M. natalensis*. For example, grooming is known to facilitate the acquisition of *Bartonella* by disrupting the skin barrier (e.g., by aggressive grooming) or by the interchange of ectoparasites between rodents (e.g., social grooming) (Gutiérrez et al., 2015). A second explanation is more in line with the main hypothesis of this study. Vanden Broecke et al. (2021a) showed that females are more likely to become infected with different helminths species and with a higher load, potentially because they consume more invertebrates (which are the secondary hosts for most of the gastrointestinal helminths) than males (Goater et al., 2014; Roberts and Janovy, 2005). The higher infection rate with those helminths might reduce the host’s immune system, making them more vulnerable to subsequent infection with *Bartonella* (Boulouis et al., 2005). Consequently, At the individual level, we found a positive co-occurrence pattern between *Bartonella* and three helminths (*P. muricola*, *T. mastomysi*, and *Trichostrongylidae*) when taking the uncertain samples into account (Fig 4 & 5). Given that we corrected for all other host characteristics and behavioral effects this may confirms the hypothesis that helminth infection lowers the Th1 immune response in *M. natalensis* and thereby facilitated co-infection with *Bartonella*.

Contrary to our expectations, we observed that less explorative individuals are more likely to become infected with *Bartonella*. This pattern has been observed previously in *M. natalensis* for both helminth and viral infections, in which less explorative individuals are more likely to be infected with different helminths and to be antibody-positive against Morogoro arenavirus (Vanden Broecke et al., 2021a, 2019). This general pattern can be explained by a potential link between exploration behavior and immunological investment, whereby highly explorative individuals are more immunological competent than less explorative individuals. Indeed, several studies have shown that bolder and more explorative individuals have a better innate immune system or more MHC alleles, which reduces the risk of infection when exposed (Garamszegi et al., 2015; Kortet et al., 2010; Zylberberg et al., 2014). An alternative explanation is that parasitic infection risk leads to behavioral modification due to sickness effects, where energy is diverted to the host immune response rather than expressing certain behaviours (Poulin, 2013).

*Anaplasma* is a genus of gram-negative intracellular, tick-borne bacteria with vertebrates as their reservoir hosts. Anaplasmosis is an important issue for farmers because the disease frequently causes abortions, reduced milk production, and increased livestock mortality (Rymaszewska and Grenda, 2008). Only the species *Anaplamsa phagocytophilum* is pathogenic for humans, as it causes human granulocytic anaplasmosis. This disease often presents as a nonspecific febrile illness, ranging from asymptomatic infections to a multi-organ failure fatal disease in immunocompromised patients (Camprubí-Ferrer et al., 2021; Jin et al., 2012). The *Anaplasma* species detected in *M. natalensis* were most closely related to uncultured *Anaplasma* (16%) and *Ehrlichia* strains (2%) previously detected in rodents and ticks from Africa. The *Anaplasma* strain was most closely related to a strain found in *Lemniscomys striatus* in Franceville in Gabon (Mangombi et al., 2021). The *Ehrlichia* strain was mainly similar to a strain detected in a *Rhipicephalus* tick (family *Ixodidae*) collected from livestock in Kisangani, Democratic Republic of Congo (Ngoy et al., 2021). Similar to *Bartonella*, we found that adult *M. natalensis* were more likely to be infected with *Anaplasma* than juveniles, which can again be explained by the long-lasting duration of infections and horizontal transmission via a vector. *Anaplasma* did not significantly co-occur with any of the helminths or *Bartonella* at the individual level. However, we observed a positive co-occurrence pattern between *Anaplasma* and *Hymenolepsis* among the different sites where they were trapped (Fig 4 & 5).This suggests that both parasites preferred a similar habitat or that similar conditions are necessary for the parasites to occur. However, it might be that this pattern results from a type I error. The prevalence of *Anaplasma* is relatively low, and we included a large number of trapping areas in our model. While this does not exclude that *Anaplasma* bacteria are spatially clustered, more data is needed to confirm this hypothesis.

*Hepatozoon* is a genus of apicomplexan protozoans that parasitize on a wide diversity of hosts (Smith, 1996). They all have a life cycle that involves an intermediate vertebrate host and a definitive blood-feeding invertebrate host (ticks, mites, sandflies, and mosquitos) (Kamani et al., 2018). Unlike most vector-borne parasites that are transmitted during a blood meal, the transmission of *Hepatozoon* to vertebrates occurs by ingestion of the infected invertebrates (Smith, 1996). Veterinarians mainly know the parasite to infect cats and dogs in which it may cause hepatozoonosis, a disease with symptoms ranging from sub-clinical in healthy pets to life-threatening in animals with extreme lethargy and anaemia (Baneth, 2011). Our screening revealed that 2.3% (5/211) of *M. natalensis* were infected with *Hepatozoon ophisauri* (99.23% similarity). This species was previously found in Iranian lizards (*Pseudopus apodus*), which are suggested to serve as an intermediate host with snakes as a final vertebrate host (Zechmeisterová et al., 2021). Besides lizards, our data indicate that rodents can be additional intermediate hosts that are prey for snakes. Indeed, predation of rodents might represent an essential route for *Hepatozoon* transmission, as rodents are more likely to eat invertebrates and snakes can be successfully infected by consuming infectious rodents’ tissues (Sloboda et al., 2008).

Overall, we found that co-infections between parasites are common and micro-macro parasite co-infections also occur, although the causal relationships underlying the patterns appear more complex than initially considered. We hypothesized that helminths would make rodents more vulnerable to microparasite infections. This is potentially true for *Bartonella* but not for *Anaplasma*. The latter also shows that the spatial distribution of different parasites can differ on a small scale, which increases the complexity of the system. Therefore, our results align with studies suggesting that interspecific interactions are less influential in structuring within-host parasite communities (Behnke et al., 2005; Haukisalmi and Henttonen, 1993; Pedersen and Fenton, 2019). One drawback of our study is that we only used cross-sectional data, limiting the ability to disentangle the effect of within-host parasite interactions from other confounding factors that can affect co-occurrence patterns (despite the use of the HMSC models that already corrected for a variety of host characteristics and spatial clustering). Longitudinal studies (e.g. capture-mark-recapture) might be better suited for this purpose as they allow insights into the causal directionality of the underlying patterns (Mariën et al., 2019b, 2018b; Pedersen and Fenton, 2019; Telfer et al., 2010).

In addition, our results confirm that *M. natalensis* can constitute multiple zoonotic pathogens of which the epidemiology is still poorly studied, especially in Africa. The fact that *M. natalensis* is often present near human dwellings suggests that humans and livestock are likely to come into contact with these parasites (Mariën et al., 2020, 2018a). While our data will allow to design further ecological studies on the risks associated to exposure with *M. natalensis*-borne parasites, medical and veterinary studies need to be conducted to elucidate the role of these parasites in human and animal pathology.

## Ethics statement

All experimental procedures were approved by the University of Antwerp Ethical Committee for Animal Experimentation (2016-63), adhered to the EEC Council Directive 2010/63/EU, and followed the Animal Ethics guidelines of the Research Policy of the Sokoine University of Agriculture.

## Acknowledgments

We thank the staff (particularly Shabani Lutea, Geoffrey Sabuni, Omary Kibwana, and Baraka Edson) at the Institute of Pest Management (Sokoine University of Agriculture, Morogoro, Tanzania) for their excellent assistance during the fieldwork. Additionally, we want to thank Pauline Van Calck and Dr. Lucinda Kirkpatrick for their supervision during the fieldwork and Dr. Joëlle Goüy de Bellocq for her incredible help during the dissections and sample preparations.

## Availability of data and materials

The biological material and data used in the current study are available from the corresponding author on reasonable request. The generated sequences were deposited in GenBank with the following accession numbers: OL982744, OL982745, OL982748 & OL984911.

## Author contributions

JM and BVB designed the study with support from HL. LB performed the fieldwork with support from BVB, LM and LM. PJJT performed the PCRs for detection of the microparasites under supervision of JM. Macroparasitological examination was done by LB and AR. BVB performed the data analysis with support from JM and HL. The first draft was written by JM and BVB. All authors contributed substantially to revisions.

## Supplementary information

**S Figure 1:**
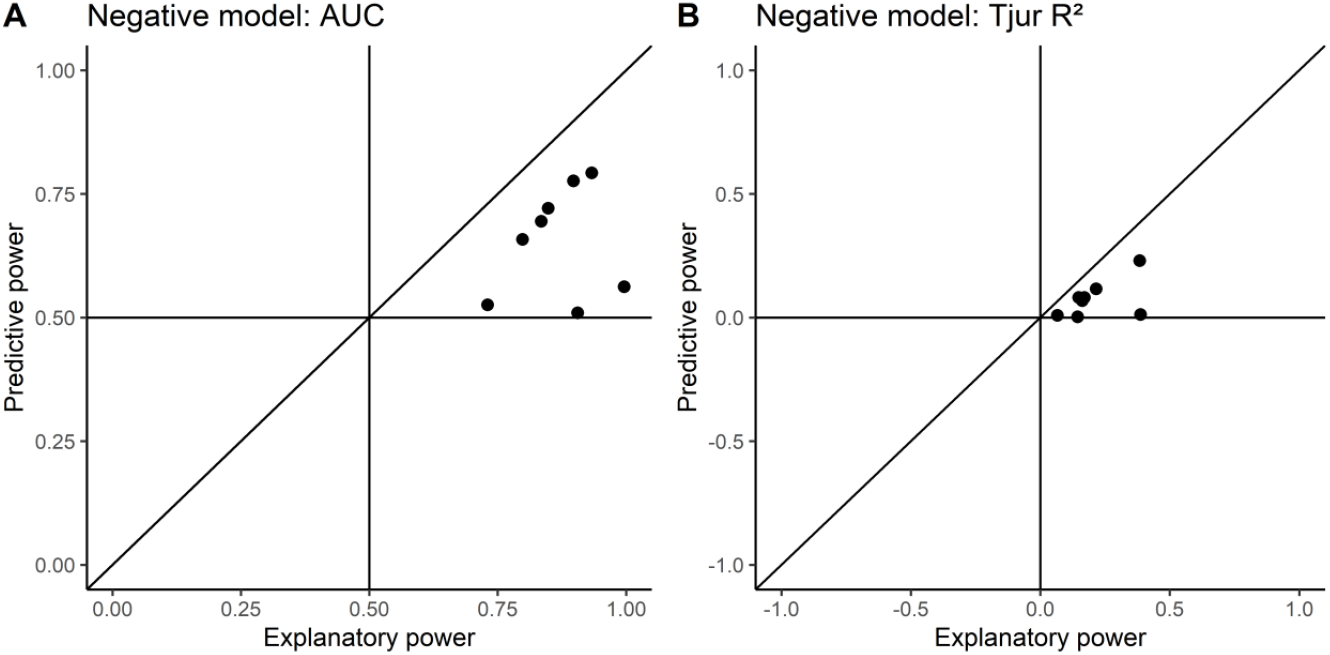
Explanatory and predictive power of the presence-absence negative model based on the A) AUC and B) Tjur R2 values, derived after five-fold cross-validation.

**S Figure 2:**
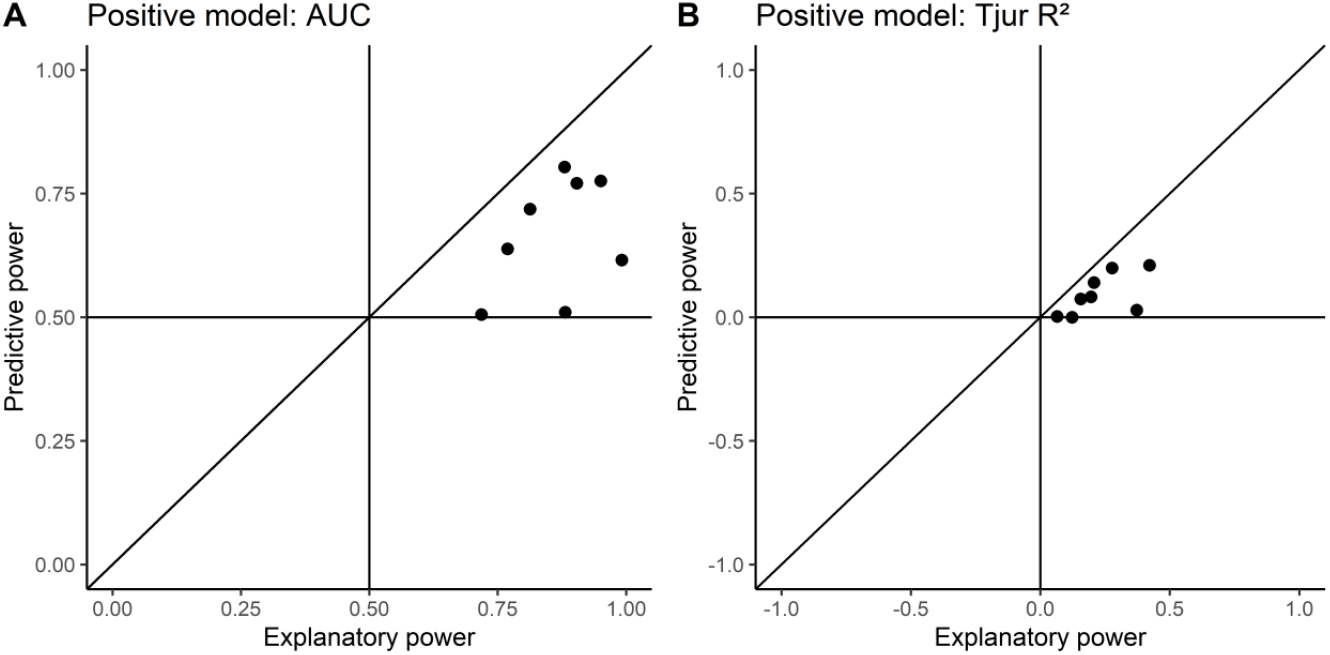
Explanatory and predictive power of the presence-absence positive model based on the A) AUC and B) Tjur R^2^ values, derived after five-fold cross-validation.

**S Figure 3:**
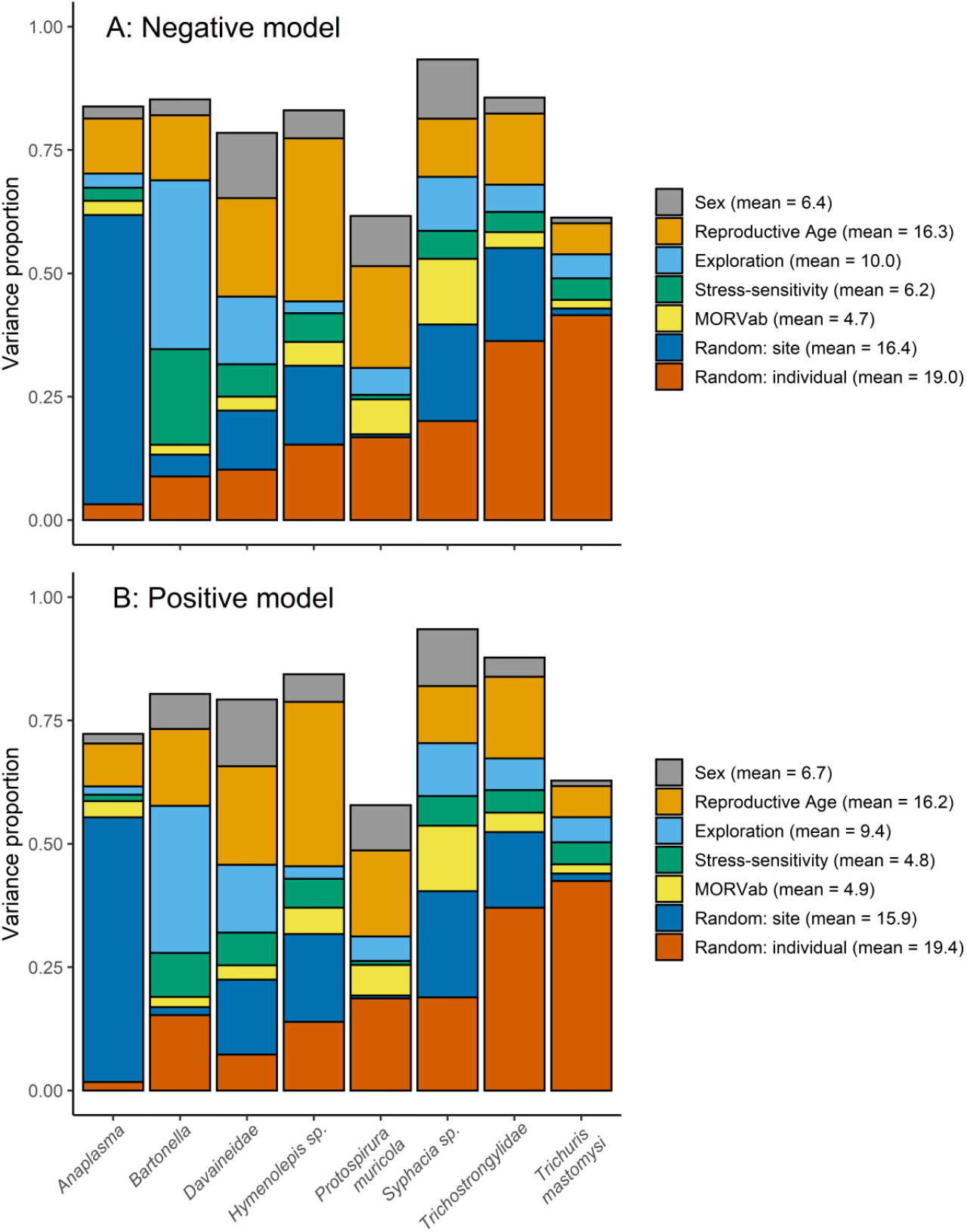
Variance partitioning among the different fixed and random effects from the (A) negative and (B) positive model. The height of the bars in both panels correspond to the explanatory power achieved by for the parasitic species measured by the Tjur R^2^. The legend gives the mean variance proportions for each fixed and random effect within the model, averaged over the species

**S table 1:** Posterior mean responses (with their 95% credibility intervals) for *Bartonella* and *Anaplasma* to the fixed effects derived from the positive model. Estimates with 95% posterior support are marked in bold.

| Fixed effect | Negative uncertain model |  | Positive uncertain model |  |
| --- | --- | --- | --- | --- |
|  | *Bartonella* | *Anaplasma* | *Bartonella* | *Anaplasma* |
|  | est (95% CrI) | est (95% CrI) | est (95% CrI) | est (95% CrI) |
| Intercept | **-1.68 (-2.20; -1.26)** | **-1.29 (-1.86; -0.81)** | **-1.41 (-1.98; -1.01)** | **-1.16 (-1.78; -0.60)** |
| Sex (male) | -0.20 (-0.79; 0.36) | 0.06 (-0.41; 0.55) | **-0.52 (-1.16; 0.04)** | -0.17 (-0.64; 0.31) |
| Age (juvenile) | **-0.75 (-1.65; 0.03)** | **-0.64 (-1.30; -0.03)** | **-0.96 (-1.89; -0.17)** | **-0.73 (-1.38; -0.11)** |
| Exploration | **-0.53 (-0.91; -0.20)** | -0.05 (-0.30; 0.19) | **-0.57 (-0.94; -0.26)** | -0.06 (-0.30; 0.18) |
| Stress-sensitivity | **-0.39 (-0.69; -0.10)** | -0.04 (-0.27; 0.20) | **-0.28 (-0.58; -0.02)** | -0.01 (-0.25; 0.22) |
| MORVab (P) | 0.03 (-0.66; 0.67) | 0.20 (-0.34; 0.75) | 0.26 (-0.36; 0.85) | 0.42 (-0.11; 0.96) |

## References

Baneth, G., 2011. Perspectives on canine and feline hepatozoonosis. Vet. Parasitol. 181, 3–11. https://doi.org/10.1016/j.vetpar.2011.04.015

Barber, I., Dingemanse, N.J., 2010. Parasitism and the evolutionary ecology of animal personality. Philos. Trans. R. Soc. Lond. B. Biol. Sci. 365, 4077–88. https://doi.org/10.1098/rstb.2010.0182

Behnke, J.M., Gilbert, F.S., Abu-Madi, M.A., Lewis, J.W., 2005. Do the helminth parasites of wood mice interact? J. Anim. Ecol. 74, 982–993. https://doi.org/10.1111/j.1365-2656.2005.00995.x

Blanco, J.L., Garcia, M.E., 2008. Immune response to fungal infections. Vet. Immunol. Immunopathol. 125, 47–70. https://doi.org/10.1016/j.vetimm.2008.04.020

Böge, I., Pfeffer, M., Htwe, N.M., Maw, P.P., Sarathchandra, S.R., Sluydts, V., Piscitelli, A.P., Jacob, J., Obiegala, A., 2021. First detection of *Bartonella* spp. In small mammals from rice storage and processing facilities in Myanmar and Sri Lanka. Microorganisms 9, 1–15. https://doi.org/10.3390/microorganisms9030658

Bohn, S.J., Webber, Q.M.R., Florko, K.R.N., Paslawski, K.R., Peterson, A.M., Piche, J.E., Menzies, A.K., Willis, C.K.R., 2017. Personality predicts ectoparasite abundance in an asocial sciurid. Ethology 123, 761–771. https://doi.org/10.1111/eth.12651

Boulouis, H.-J., Chao-chin, C., Henn, J.B., Kasten, R.W., Chomel, B.B., 2005. Factors associated with the rapid emergence of zoonotic *Bartonella* infections. Vet. Res. 36, 383–410. https://doi.org/10.1051/vetres:2005009

Boyer, N., Réale, D., Marmet, J., Pisanu, B., Chapuis, J.-L., 2010. Personality, space use and tick load in an introduced population of Siberian chipmunks *Tamias sibiricus.* J. Anim. Ecol. 79, 538–547. https://doi.org/10.1111/j.1365-2656.2010.01659.x

Brouat, C., Duplantier, J.-M., 2007. Host habitat patchiness and the distance decay of similarity among gastro-intestinal nematode communities in two species of *Mastomys* (southeastern Senegal). Oecologia 152, 715–720. https://doi.org/10.1007/s00442-007-0680-8

Brouat, C., Kane, M., Diouf, M., Bâ, K., Sall-Dramé, R., Duplantier, J.M., 2007. Host ecology and variation in helminth community structure in *Mastomys* rodents from Senegal. Parasitology 134, 437–450. https://doi.org/10.1017/S003118200600151X

Camprubí-Ferrer, D., Portillo, A., Santibáñez, S., Almuedo-Riera, A., Rodriguez-Valero, N., Subirà, C., Martinez, M.J., Navero-Castillejos, J., Fernandez-Pardos, M., Genton, B., Cobuccio, L., Van Den Broucke, S., Bottieau, E., Muñoz, J., Oteo, J.A., 2021. Incidence of human granulocytic anaplasmosis in returning travellers with fever. J. Travel Med. 28, 1–6. https://doi.org/10.1093/jtm/taab056

Clark, N.J., Wells, K., Dimitrov, D., Clegg, S.M., 2016. Co-infections and environmental conditions drive the distributions of blood parasites in wild birds. J. Anim. Ecol. 85, 1461–1470. https://doi.org/10.1111/1365-2656.12578

Dahmana, H., Granjon, L., Diagne, C., Davoust, B., Fenollar, F., Mediannikov, O., 2020. Rodents as hosts of pathogens and related zoonotic disease risk. Pathogens 9. https://doi.org/10.3390/pathogens9030202

Dahmani, M., Diatta, G., Labas, N., Diop, A., Bassene, H., Raoult, D., Granjon, L., Fenollar, F., Mediannikov, O., 2018. Noncontiguous finished genome sequence and description of *Bartonella mastomydis* sp. nov. New Microbes New Infect. 25, 60–70. https://doi.org/10.1016/j.nmni.2018.03.005

Dehio, C., 2004. Molecular and cellular basis of *bartonella* pathogenesis. Annu. Rev. Microbiol. 58, 365–390. https://doi.org/10.1146/annurev.micro.58.030603.123700

Diagne, C., Ribas, A., Charbonnel, N., Dalecky, A., Tatard, C., Gauthier, P., Haukisalmi, V., Fossati-Gaschignard, O., Bâ, K., Kane, M., Niang, Y., Diallo, M., Sow, A., Piry, S., Sembène, M., Brouat, C., 2016. Parasites and invasions: changes in gastrointestinal helminth assemblages in invasive and native rodents in Senegal. Int. J. Parasitol. 46, 857–869. https://doi.org/10.1016/j.ijpara.2016.07.007

Diouf, M., Diagne, C.A., Quilichini, Y., Dobigny, G., Garba, M., Marchand, B., 2013. *Pterygodermatites (Mesopectines) niameyensis* n. sp. (Nematoda: Rictulariidae), a parasite of *Mastomys natalensis* (Smith, 1834) (Rodentia: Muridae) from Niger. J. Parasitol. 99, 1034–1039. https://doi.org/10.1645/13-204.1

Ezenwa, V.O., Etienne, R.S., Luikart, G., Beja-Pereira, A., Jolles, A.E., 2010. Hidden consequences of living in a wormy world: Nematode-induced immune suppression facilitates tuberculosis invasion in African buffalo. Am. Nat. 176, 613–624. https://doi.org/10.1086/656496

Garamszegi, L.Z., Zagalska-Neubauer, M., Canal, D., Markó, G., Szász, E., Zsebők, S., Szöllősi, E., Herczeg, G., Török, J., 2015. Malaria parasites, immune challenge, MHC variability, and predator avoidance in a passerine bird. Behav. Ecol. 26, 1292–1302. https://doi.org/10.1093/beheco/arv077

Goater, T.M., Goater, C.P., Esch, G.W., 2014. Parasitism: The Diversity and Ecology of Animal Parasites, Second edi. ed. Cambridge University Press, New York. https://doi.org/10.1080/15627020.2002.11657185

Gutiérrez, R., Krasnov, B., Morick, D., Gottlieb, Y., Khokhlova, I.S., Harrus, S., 2015. *Bartonella* infection in rodents and their flea ectoparasites: An overview. Vector-Borne Zoonotic Dis. 15, 27–39. https://doi.org/10.1089/vbz.2014.1606

Haukisalmi, V., Henttonen, H., 1993. Coexistence in helminths of the bank vole Clethrionomys glareolus. I. Patterns of co-occurrence. J. Anim. Ecol. 62, 221–229.

Henrichs, B., Oosthuizen, M.C., Troskie, M., Gorsich, E., Gondhalekar, C., Beechler, B.R., Ezenwa, V.O., Jolles, A.E., 2016. Within guild co-infections influence parasite community membership: a longitudinal study in African Buffalo. J. Anim. Ecol. 85, 1025–1034. https://doi.org/10.1111/1365-2656.12535

Holt, J., Davis, S., Leirs, H., 2006. A model of Leptospirosis infection in an African rodent to determine risk to humans: Seasonal fluctuations and the impact of rodent control. Acta Trop. 99, 218–225. https://doi.org/10.1016/j.actatropica.2006.08.003

Jin, H., Wei, F., Liu, Q., Qian, J., 2012. Epidemiology and control of human granulocytic anaplasmosis: A systematic review. Vector-Borne Zoonotic Dis. 12, 269–274. https://doi.org/10.1089/vbz.2011.0753

Kamani, J., Harrus, S., Nachum-Biala, Y., Gutiérrez, R., Mumcuoglu, K.Y., Baneth, G., 2018. Prevalence of *Hepatozoon* and *Sarcocystis* spp. in rodents and their ectoparasites in Nigeria. Acta Trop. 187, 124–128. https://doi.org/10.1016/j.actatropica.2018.07.028

Kortet, R., Hedrick, A. V, Vainikka, A., 2010. Parasitism, predation and the evolution of animal personalities. Ecol. Lett. 13, 1449–1458. https://doi.org/10.1111/j.1461-0248.2010.01536.x

Leirs, H., Verhagen, R., Verheyen, W., 1994. The basis of reproductive seasonality in *Mastomys* rats (Rodentia: Muridae) in Tanzania. J. Trop. Ecol. 10, 55–66.

Leirs, H., Verheyen, W., Michiels, M., Verhagen, R., Stuyck, J., 1990. The relation between rainfall and the breeding season of mastomys natalensis (smith, 1834) in morogoro, tanzania. Ann. la société R. Zool. Belgique 119, 59–64.

Mangombi, J.B., N’dilimabaka, N., Lekana-Douki, J.-B., Banga, O., Maghendji-nzondo, S., Id, M.B., Leroy, E., Fenollar, F., Mediannikov, O., 2021. First investigation of pathogenic bacteria, protozoa and viruses in rodents and shrews in context of forest-savannah-urban areas interface in the city of Franceville (Gabon). PLoS One 1–28. https://doi.org/10.1371/journal.pone.0248244

Mariën, J., Borremans, B., Kourouma, F., Baforday, J., Rieger, T., Günther, S., Magassouba, N., Leirs, H., Fichet-Calvet, E., 2019a. Evaluation of rodent control to fight Lassa fever based on field data and mathematical modelling. Emerg. Microbes Infect. 8, 640–649. https://doi.org/10.1080/22221751.2019.1605846

Mariën, J., Borremans, B., Verhaeren, C., Kirkpatrick, L., Gryseels, S., Goüy de Bellocq, J., Günther, S., Sabuni, C.A., Massawe, A.W., Reijniers, J., Leirs, H., 2019b. Density dependence and persistence of Morogoro arenavirus transmission in a fluctuating population of its reservoir host. J. Anim. Ecol. 89, 1365–2656. https://doi.org/10.1111/1365-2656.13107

Mariën, J., Iacono, G. Lo, Rieger, T., Magassouba, N., Günther, S., Fichet-calvet, E., Mariën, J., Iacono, G. Lo, Rieger, T., Magassouba, N., 2020. Households as hotspots of Lassa fever ? Assessing the spatial distribution of Lassa virus-infected rodents in rural villages of Guinea. https://doi.org/10.1080/22221751.2020.1766381

Mariën, J., Kourouma, F., Magassouba, N., Leirs, H., Fichet-Calvet, E., 2018a. Movement patterns of small rodents in Lassa fever-endemic villages in Guinea. Ecohealth 15, 348–359. https://doi.org/10.1007/s10393-018-1331-8

Mariën, J., Sluydts, V., Borremans, B., Gryseels, S., Vanden Broecke, B., Sabuni, C.A., Katakweba, A.A.S., Mulungu, L.S., Günther, S., de Bellocq, J.G., Massawe, A.W., Leirs, H., 2018b. Arenavirus infection correlates with lower survival of its natural rodent host in a long-term capture-mark-recapture study. Parasit. Vectors 11, 90. https://doi.org/10.1186/s13071-018-2674-2

McArdle, A.J., Turkova, A., Cunnington, A.J., 2018. When do co-infections matter? Curr. Opin. Infect. Dis. 31, 209–215. https://doi.org/10.1097/QCO.0000000000000447

Meerburg, B.G., Singleton, G.R., Kijlstra, A., 2009. Rodent-borne diseases and their risks for public health, Critical Reviews in Microbiology. 35, 221–70. https://doi.org/10.1080/10408410902989837

Monath, T.P., 1987. Lassa fever: new issues raised by field studies in West Africa. J. Infect. Dis. 155, 433–436. https://doi.org/10.1093/infdis/155.3.433

Mulungu, L.S., 2017. Control of rodent pests in maize cultivation: the case of Africa 317–337. https://doi.org/10.19103/AS.2016.0002.18

Mwanjabe, P.S., Sirima, F.B., Lusingu, J., 2002. Crop losses due to outbreaks of *Mastomys natalensis* (Smith, 1834) Muridae, Rodentia, in the Lindi Region of Tanzania. Int. Biodeterior. Biodegradation 49, 133–137. https://doi.org/10.1016/S0964-8305(01)00113-5

Neerinckx, S.B., Peterson, A.T., Gulinck, H., Deckers, J., Leirs, H., 2008. Geographic distribution and ecological niche of plague in sub-Saharan Africa. Int. J. Health Geogr. 7, 54. https://doi.org/10.1186/1476-072X-7-54

Ngoy, S., Diarra, A.Z., Laudisoit, A., Gembu, G.C., Verheyen, E., Mubenga, O., Mbalitini, S.G., Baelo, P., Laroche, M., Parola, P., 2021. Using MALDI-TOF mass spectrometry to identify ticks collected on domestic and wild animals from the Democratic Republic of the Congo. Exp. Appl. Acarol. 84, 637–657. https://doi.org/10.1007/s10493-021-00629-z

Oguge, N., Rarieya, M., Ondiaka, P., 1997. A preliminary survey of macroparasite communities of rodents of Kahawa, central Kenya. Belgian J. Zool. 127, 113–118.

Ovaskainen, O., Abrego, N., 2020. Joint Species Distribution Modelling. Cambridge University Press, Cambridge. https://doi.org/10.1017/9781108591720

Ovaskainen, O., Tikhonov, G., Norberg, A., Guillaume Blanchet, F., Duan, L., Dunson, D., Roslin, T., Abrego, N., 2017. How to make more out of community data? A conceptual framework and its implementation as models and software. Ecol. Lett. 20, 561–576. https://doi.org/10.1111/ele.12757

Pearce, J., Ferrier, S., 2000. Evaluating the predictive performance of habitat models developed using logistic regression. Ecol. Modell. 133, 225–245. https://doi.org/10.1016/S0304-3800(00)00322-7

Pedersen, A.B., Fenton, A., 2019. Wild rodents as a natural model to study within-host parasite interactions, in: Wildlife Disease Ecology. pp. 58–90. https://doi.org/10.1017/9781316479964.003

Poulin, R., 2013. Parasite manipulation of host personality and behavioural syndromes. J. Exp. Biol. 216, 18–26. https://doi.org/10.1242/jeb.073353

Ramsay, C., Rohr, J.R., 2021. The application of community ecology theory to co-infections in wildlife hosts. Ecology 102, 0–2. https://doi.org/10.1002/ecy.3253

Ribas, A., Chaisiri, K., Morand, S., Hugot, J.-P., Haukisalmi, V., Henttonen, H., 2011. Isolating helminths in rodents, in: Herbreteau, V., Jittapalapong, S., Rerkamnuaychoke, W., Chaval, Y., Cosson, J.-F., Morand, S. (Eds.), Protocols for Field and Laboratory Rodent Studies. Kasetsart University Press. https://doi.org/10.13140/2.1.2353.4406

Ribas, A., Diagne, C., Tatard, C., Diallo, M., Poonlaphdecha, S., Brouat, C., 2017. Whipworm diversity in West African rodents: a molecular approach and the description of *Trichuris duplantieri* n. sp. (Nematoda: Trichuridae). Parasitol. Res. 116, 1265–1271. https://doi.org/10.1007/s00436-017-5404-3

Ribas, A., López, S., Makundi, R.H., Leirs, H., de Bellocq, J.G., 2013. Trichuris spp. (Nematoda: *Trichuridae*) from two rodents, *Mastomys natalensis* and *Gerbilliscus vicinus* in Tanzania. J. Parasitol. 99, 868–75. https://doi.org/10.1645/12-151.1

Ribas, A., Makundi, R.H., Bellocq, J.G. De, 2012. *Paraconcinnum leirsi* n.sp. (Trematoda: *Dicrocoeliidae)* from Rodents in Tanzania and its Phylogenetic Position within the *Dicrocoeliids.* African Zool. 47, 326–331. https://doi.org/10.3377/004.047.0219

Roberts, L.S., Janovy, J.J., 2005. Foundations of Parasitology, Seventh ed. ed. McGraw-Hill, New York.

Rymaszewska, a., Grenda, S., 2008. Anaplasma–characteristics of *Anaplasma* and their vectors: a review. Vet. Med. (Praha). 11, 573–584.

Sadlova, J., Vojtkova, B., Hrncirova, K., Lestinova, T., Spitzova, T., Becvar, T., Votypka, J., Bates, P., Volf, P., 2019. Host competence of African rodents Arvicanthis neumanni, A. niloticus and *Mastomys natalensis* for Leishmania major. Int. J. Parasitol. Parasites Wildl. 8, 118–126. https://doi.org/10.1016/j.ijppaw.2019.01.004

Salvador, A.R., Guivier, E., Xuéreb, A., Chaval, Y., Cadet, P., Poulle, M.L., Sironen, T., Voutilainen, L., Henttonen, H., Cosson, J.F., Charbonnel, N., 2011. Concomitant influence of helminth infection and landscape on the distribution of Puumala hantavirus in its reservoir, *Myodes glareolus.* BMC Microbiol. 11, 30. https://doi.org/10.1186/1471-2180-11-30

Sloboda, M., Kamler, M., Bulantová, J., Votýpka, J., Modrý, D., 2008. Rodents as intermediate hosts of *Hepatozoon ayorgbor* (Apicomplexa: Adeleina: *Hepatozoidae)* from the African ball python, *Python regius?* Folia Parasitol. (Praha). 55, 13–16. https://doi.org/10.14411/fp.2008.003

Smith, T.G., 1996. The genus *Hepatozoon* (Apicomplexa: *Adeleina).* J. Parasitol. 82, 565–585. https://doi.org/10.2307/3283781

Telfer, S., Lambin, X., Birtles, R., Beldomenico, P., Burthe, S., Paterson, S., Begon, M., 2010. Species interactions in a parasite community drive infection risk in a wildlife population. Science (80-.). 330, 243–246. https://doi.org/10.1126/science.1190333

Tikhonov, G., Opedal, Ø.H., Abrego, N., Lehikoinen, A., de Jonge, M.M.J., Oksanen, J., Ovaskainen, O., 2020. Joint species distribution modelling with the r-package Hmsc. Methods Ecol. Evol. 11, 442–447. https://doi.org/10.1111/2041-210X.13345

Tjur, T., 2009. Coefficients of determination in logistic regression models - A new proposal: The coefficient of discrimination. Am. Stat. 63, 366–372. https://doi.org/10.1198/tast.2009.08210

Vanden Broecke, B., Bernaerts, L., Ribas, A., Sluydts, V., Mnyone, L., Matthysen, E., Leirs, H., 2021a. Linking behavior, co-infection patterns, and viral infection risk with the whole gastrointestinal helminth community structure in *Mastomys natalensis.* Front. Vet. Sci. 8, 1–15. https://doi.org/10.3389/fvets.2021.669058

Vanden Broecke, B., Bongers, A., Mnyone, L., Matthysen, E., Leirs, H., 2021b. Nonlinear maternal effects on personality in a rodent species with fluctuating densities. Curr. Zool. 67, 1–9. https://doi.org/10.1093/cz/zoaa032

Vanden Broecke, B., Mariën, J., Sabuni, C.A., Mnyone, L., Massawe, A.W., Matthysen, E., Leirs, H., 2019. Relationship between population density and viral infection: A role for personality? Ecol. Evol. 9, 10213–10224. https://doi.org/10.1002/ece3.5541

Vanden Broecke, B., Sluydts, V., Mariën, J., Sabuni, C.A., Massawe, A.W., Matthysen, E., Leirs, H., 2021c. The effects of personality on survival and trappability in a wild mouse during a population cycle. Oecologia 195, 901–913. https://doi.org/10.1007/s00442-021-04897-9

Vaumourin, E., Vourc’h, G., Gasqui, P., Vayssier-Taussat, M., 2015. The importance of multiparasitism: examining the consequences of co-infections for human and animal health. Parasit. Vectors 8, 545. https://doi.org/10.1186/s13071-015-1167-9

Welc-Faleciak, R., Bajer, A., Behnke, J.M., Siński, E., 2010. The ecology of *Bartonella* spp. infections in two rodent communities in the Mazury Lake District region of Poland. Parasitology 137, 1069–1077. https://doi.org/10.1017/S0031182009992058

Zechmeisterová, K., Javanbakht, H., Kvičerová, J., Široký, P., 2021. Against growing synonymy: Identification pitfalls of *Hepatozoon* and *Schellackia* demonstrated on North Iranian reptiles. Eur. J. Protistol. 79, 1–18. https://doi.org/10.1016/j.ejop.2021.125780

Zylberberg, M., Klasing, K.C., Hahn, T.P., 2014. In house finches, Haemorhous mexicanus, risk takers invest more in innate immune function. Anim. Behav. 89, 115–122. https://doi.org/10.1016/j.anbehav.2013.12.021

